## Extended Data for "SpatialDM: Rapid identification of spatially co-expressed ligand-receptor reveals cell-cell communication patterns"

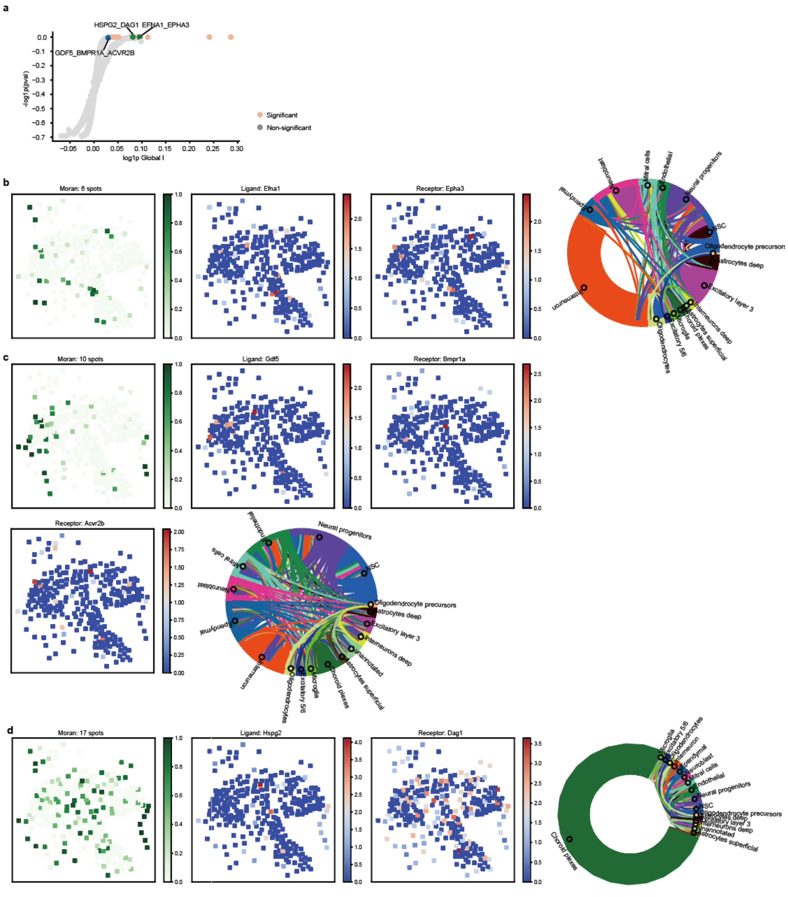

Fig. 1 Detecting spatial LRI in mouse ventricular-subventricular zone (SVZ) We demonstrated the broad applicability of SpatialDM from Next-Generation Sequencing data to Fluorescent In Situ Hybridization (FISH) data. SVZ, located along the walls of the brain lateral ventricles, is the birthplace for neural stem cells throughout life. Many LRIs have been identified essential to the neurogenic niche, including mitogenic signals like fibroblast growth factor 2 (FGF-2) and epidermal growth factor (EGF), neurogenic signals like BMP and Shh, as well as membrane-bound signals like Notch and Eph [1, 2]. (a) From the limited insights due to low sequencing coverage of FISH-based sequencing, we found (b) EFNA1_EPHA3 enriched in neural progenitors, neuroblast and neural NSCs [2], (c) GDF5 signaling to BMPR1A_ACVR2B complex from various cell types, in particular NSC and neural progenitors [3], and (d) HSPG2_DAG1 transmitted between adjacent choroid plexes cells [4], etc. In the Moran p-value spatial plots, dot color indicates 1 - p-values (of all ranges), the selected number of spots refers to only spots with the uncorrected $p < 0.1$. The chord diagram visualizes the dominant cell types where edge numbers correspond to weighted cell type composition on selected spots and the edge color indicates sender cell types.

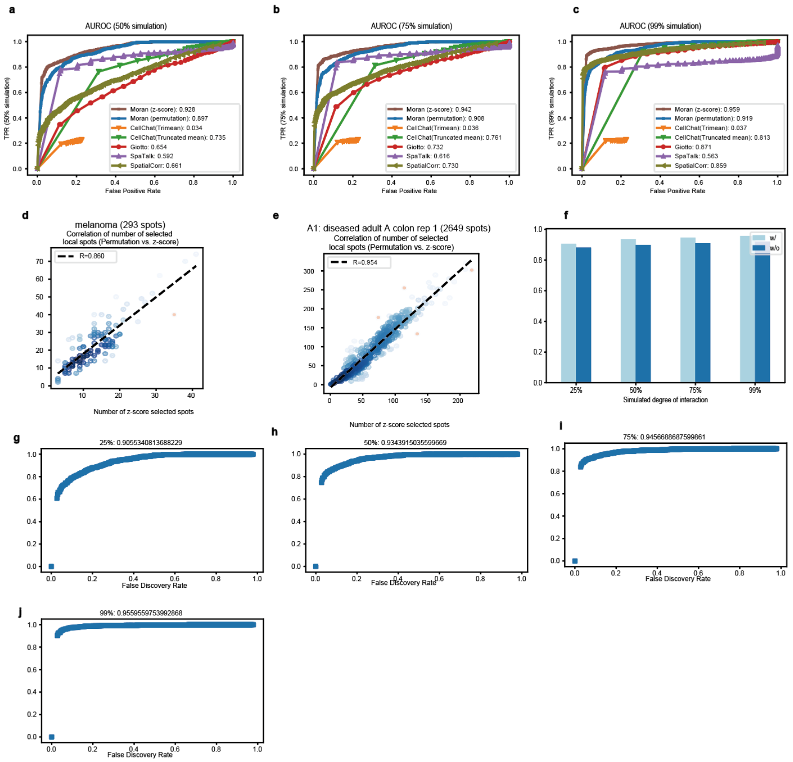


Fig. 2 (a-c) ROC under the 50% (a), 75% (b), 99% (c) degrees of interaction simulation scenario. AUROC for each method is labeled in the legend. (c-d) Correlations of the numbers of local selected spots ( $p < 0.1$) between z-score approach and permutation approach, in the melanoma dataset (d) and the intestine dataset (e). (f) AUROC comparison for global Moran permutation tests with (0.17 weight) vs. without considering ligand auto-correlation in each simulation scenario. (g-j) ROC under the 50% (a), 75% (b), 99% (c) degrees of interaction after considering ligand auto-correlation.

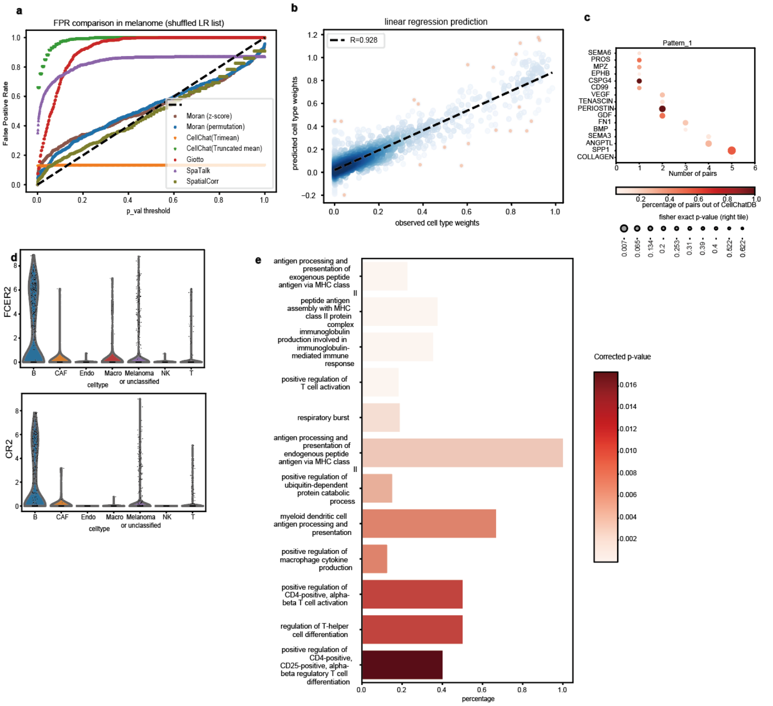


Fig. 3 (a) FPR comparison of SpatialDM with CellChat, Giotto, SpaTalk, and SpatialCorr in the melanoma dataset but with shuffled LR pairs. Only SpatialDM (both z-score and permutation approaches) calibrates with the null distribution along with the diagonal line. (b) Cell type prediction based on local Moran p-values. The whole dataset was used for fitting the linear regression model and for prediction. An Pearson coefficient R = 0 .928 was observed. (c) Pathway enrichment result for pattern 3 interactions. (d) FCER2 and CR2 expression across different cell types in the scRNA dataset. (e) Gene Ontology enrichment of the 3500 genes that are up-regulated in the CD23 hot spots.

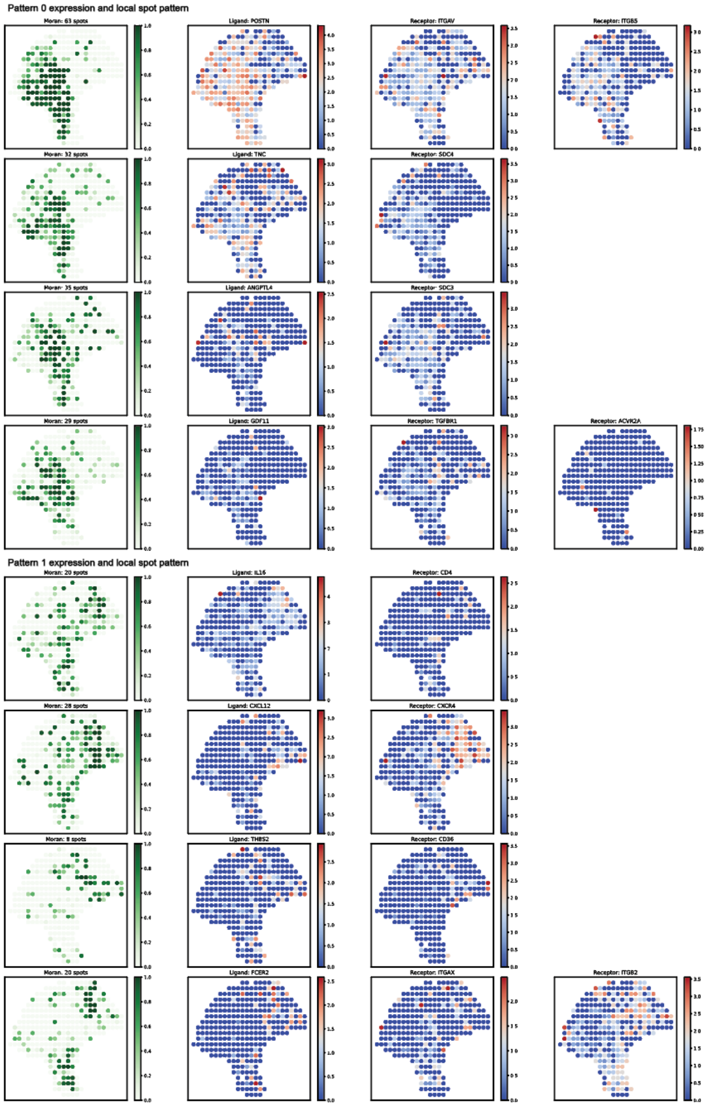


Fig. 4 For 4 exemplar interactions belonging to pattern 0 (Upper panel) and pattern 1 (Lower panel), respectively, the local Moran spots (permutation, $p < 0.1$), and their corresponding LR expression were visualized. Local p of all ranges are displayed.

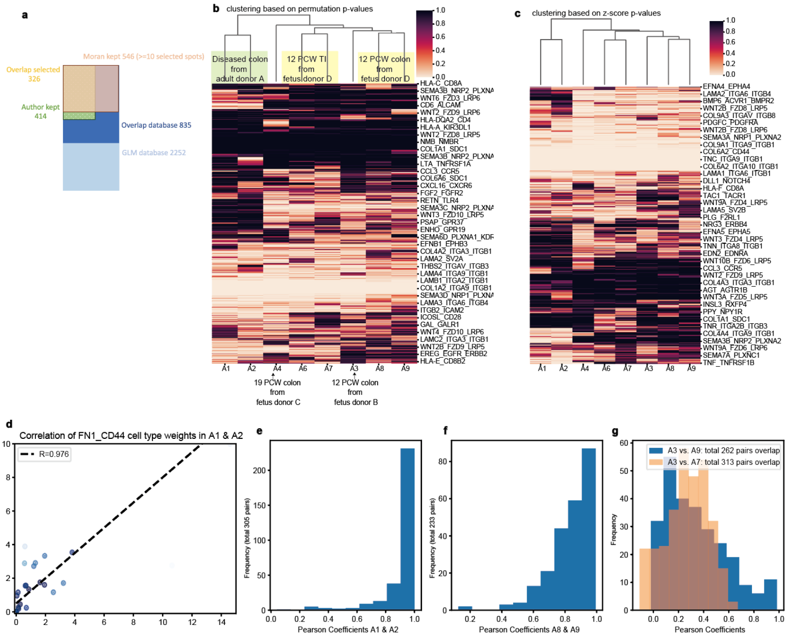


Fig. 5 (a) Comparison of the number of selected pairs across all 8 samples by Corbett, et al. vs. global Moran (permutation approach, $p < 0.05$). (b-c) Cluster map based on permutaion (b) or z-score (c) p-values. Each row is an interaction; each column is a sample. The sample clustering results were visualized on the top, which were consistent with sample kniship. (d) For FN1 _CD44, the correlation of additive cell type weights across selected spots between A1 and A2 (permutation, $p < 0.1$). Each dot represents a cell type. x- and y-axis represent additive cell type weights. An Pearson coefficient $R = 0.991$ was observed. (e-g) Summary coefficient histogram for 4 pairs of samples (i.e. Technical replicates A1 vs. A2 in E, A8 vs. A9 in F, biological replicates A3 vs. A9 and non-biological replicates A3 vs. A7 in g). Each statistic was a Pearson coefficient computed for an overlapping pair between the samples. The total number of overlapping pairs were specified in y-axis labels or legend.

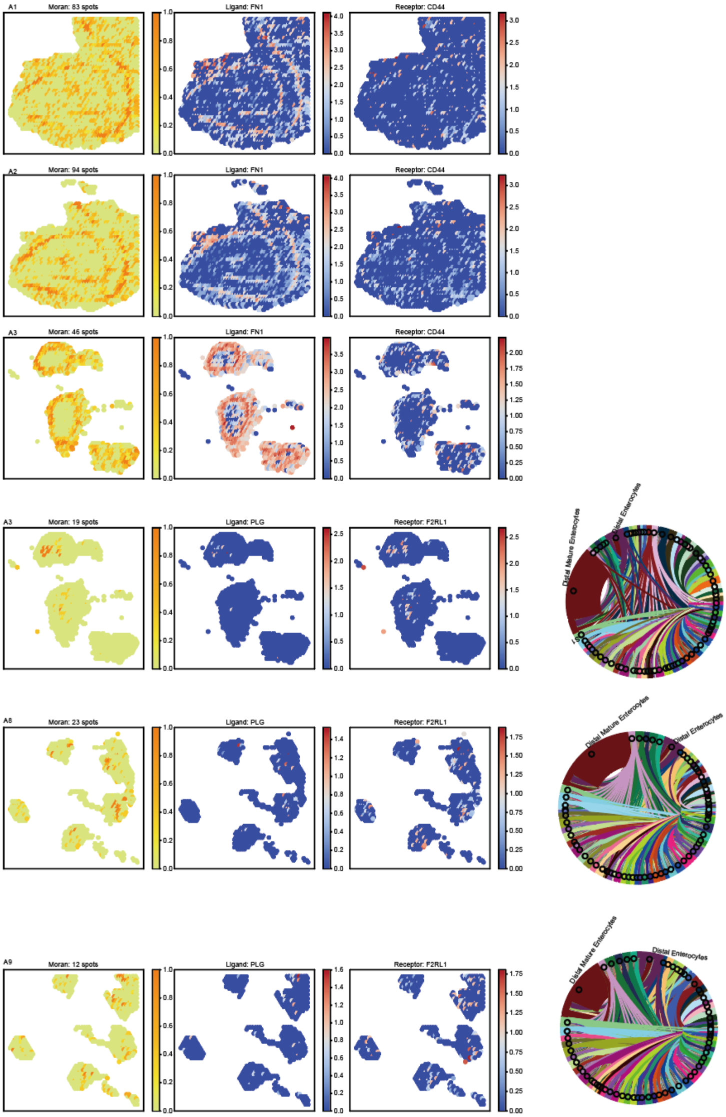


Fig. 6 Spatial plots of FN1 _CD44 (Upper) and PLG _F2RL1 (Lower) in the specified samples. For each lane, local Moran selected spots (z-score approach, colored by the value of 1 − p), LR expression, and (for PLG _F2RL1) the cell type across selected spots were visualized. In the Moran p-value spatial plots, dot color indicates 1 - p-values (of all ranges), the selected number of spots refers to only spots with the uncorrected $p < 0.1$. The chord diagram visualizes the dominant cell types where edge numbers correspond to weighted cell type composition on selected spots and the edge color indicates sender cell types.

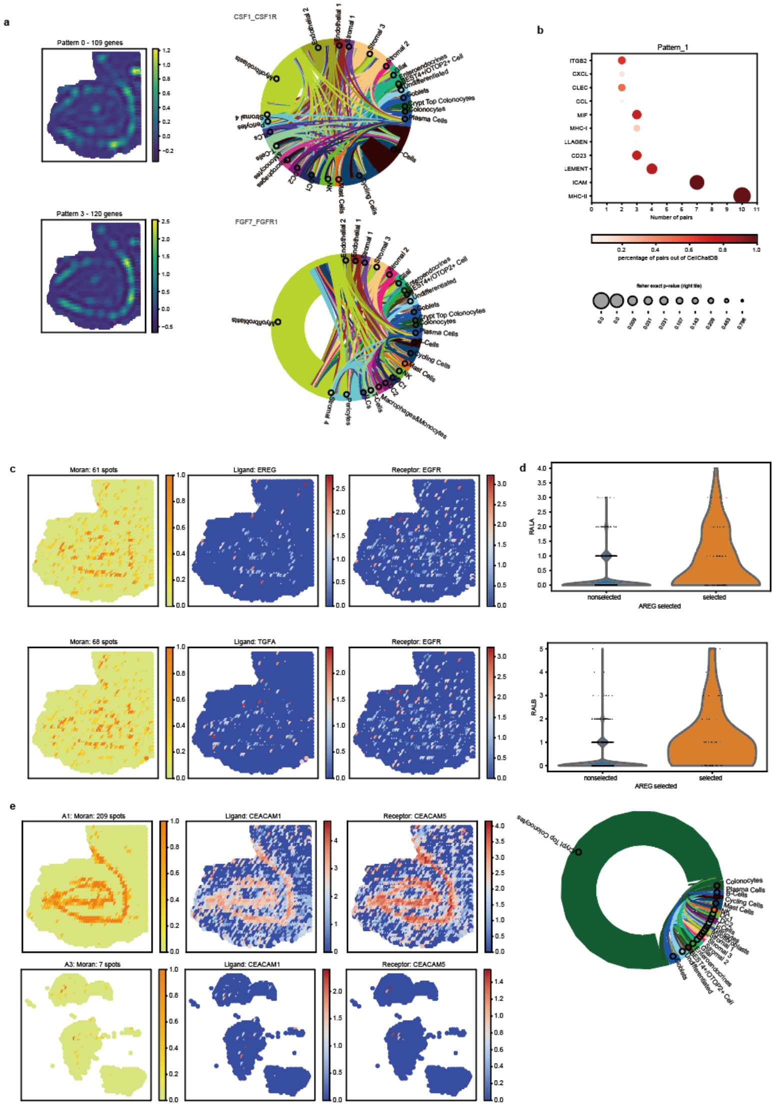

Fig. 7 (a) Similar to Fig. 3D, pattern 2 and 3 summary for A1 SpatialDE results. (b) Pathway enrichment results for pattern 0 and 2. (c) Similar to Supp Fig 6, two other EGF pathway pairs were visualized in A1, in addition to Fig. 3F. (d) Violin plots of EGF down-stream gene expression. (e) The CEACAM1 _CEACAM5 interaction pattern visualized in A1 (ubiquitous) versus A3 (sparse).

### References

[1] Lim, D.A., Alvarez-Buylla, A.: The adult ventricular–subventricular zone (v-svz) and olfactory bulb (ob) neurogenesis. Cold Spring Harbor perspectives in biology 8(5), 018820 (2016)

[2] Tong, C.K., Alvarez-Buylla, A.: Snapshot: adult neurogenesis in the v-svz. Neuron 81(1), 220 (2014)

[3] Marcy, G., Foucault, L., Babina, E., Texeraud, E., Zweifel, S., Heinrich, C., Hernandez-Vargas, H., Parras, C., Jabaudon, D., Raineteau, O.: Single cell analysis of the dorsal v-svz reveals differential quiescence of postnatal pallial and sub-pallial neural stem cells driven by tgfbeta/bmp-signalling. bioRxiv (2022) https://arxiv.org/abs/https://www.biorxiv.org/content/early/2022/05/20/2022.05.20.492790.full.pdf . https://doi.org/10.1101/2022.05.20.492790

[4] Lun, M.P., Johnson, M.B., Broadbelt, K.G., Watanabe, M., Kang, Y.-j., Chau, K.F., Springel, M.W., Malesz, A., Sousa, A.M., Pletikos, M., et al.: Spatially heterogeneous choroid plexus transcriptomes encode positional identity and contribute to regional csf production. Journal of Neuroscience 35(12), 4903–4916 (2015)
