## Supplementary material for "SpatialDM: Rapid identification of spatially co-expressed ligand-receptor reveals cell-cell communication patterns": Supp. Note 1

### Supplementary Notes of SpatialDM

#### 1 Supplementary Note 1: analytical null distribution of bi-variate Moran's $R$

##### 1.1 Univariate Moran's $I$

Univariate Moran's $I$ is established for hypothesis testing on the spatial autocorrelation of a single variable $\boldsymbol{x}$. Let $\boldsymbol{x} = (x_1, x_2, ..., x_n)$ be the vector of values over $n$ locations, and $\bar{x}$ be the mean of $\boldsymbol{x}$. $W = w_{[ij]}$ is the spatial weight matrix (without normalization). The formula of the statistic takes the form in the following equation:

$$I = \frac{n}{\sum_{i=1}^{n}\sum_{j=1}^{n} w_{ij}} \frac{\sum_{i=1}^{n}\sum_{j=1}^{n} w_{ij}(x_i - \bar{x})(x_j - \bar{x})}{\sum_{i=1}^{n}(x_i - \bar{x})^2}$$

Null hypothesis assumes that

1) $x$ is normally distributed
2) $x$ is spatially randomly distributed

Under the assumption, the expectation and variance of univariate Moran's $I$ are

$$E(I) = \frac{-1}{n-1}$$

$$Var(I) = \frac{n((n^2 - 3n + 3)S_1 - nS_2 + 3(\sum_{i=1}^{n}\sum_{j=1}^{n} w_{ij})^2}{(n-1)(n-2)(n-3)(\sum_{i=1}^{n}\sum_{j=1}^{n} w_{ij})^2} - (E(I))^2$$

$$S_1 = \frac{1}{2}\sum_{i=1}^{n}\sum_{j=1}^{n}(w_{ij} + w_{ji})^2, \quad S_2 = \sum_{i=1}^{n}(\sum_{j=1}^{n} w_{ij} + \sum_{j=1}^{n} w_{ji})^2$$

##### 1.2 Bivariate Moran's global $R$

Here, in order to distinguish the univariate auto-correlation, we use the symbol $R$ to indicate spatial correlation. Specifically, the multivariate Moran's $R$ indicates the spatial correlation between one variable and another variable in neighbouring regions. Here we consider the bivariate case. Let $\boldsymbol{X} = (X_1, X_2, ..., X_n)$ and $\boldsymbol{Y} = (Y_1, Y_2, ..., Y_n)$ be the two variables of interest. The formula of the statistic takes the form in the following equation:

$$R = \frac{n}{\sum_{i=1}^{n}\sum_{j=1}^{n} w_{i,j}} \frac{\sum_{i=1}^{n}\sum_{j=1}^{n} w_{ij}(X_i - \bar{X})(Y_j - \bar{Y})}{\sqrt{\sum_{i=1}^{n}(X_i - \bar{X})^2}\sqrt{\sum_{i=1}^{n}(Y_i - \bar{Y})^2}} \tag{1}$$

The null hypothesis assumes that

1) $X$ and $Y$ are normally distributed: $X \sim N(0, \sigma_1^2 I), Y \sim N(0, \sigma_2^2 I)$.

2) $X$ and $Y$ are is spatially randomly distributed.

The expectation and variance of the test statistic need to be derived under the null hypothesis to conduct the z-score test. Note, in Eq. (1), we can further absorb the normalisation term of $W$ to simplify the notation. However, the de-mean operations over $X$ and $Y$ cannot be absorbed for simplification, as it will break the i.i.d. assumption. Instead, we have to introduce a centering matrix below.

Denote $H$ as the centering matrix, $H = I - \frac{1}{n}\mathbf{1}\mathbf{1}^T$, then Eq. (1) can be rewritten in matrix form:

$$\begin{aligned} R &= \frac{n}{\sum_{i=1}^{n}\sum_{j=1}^{n} w_{i,j}} \frac{(HX)^T WHY}{\sqrt{(HX)^T HX}\sqrt{(HY)^T HY}} \\ &= \frac{n}{\sum_{i=1}^{n}\sum_{j=1}^{n} w_{i,j}} \frac{X^T HWHY}{\sqrt{X^T HHX}\sqrt{Y^T HHY}} \\ &= \frac{n}{\sum_{i=1}^{n}\sum_{j=1}^{n} w_{i,j}} \frac{X^T HWHY}{\sqrt{X^T HXY^T HY}} \end{aligned} \tag{2}$$

Since $H$ is symmetric, diagonalizing $H$ will give

$$H = P^T Diag(1, 1, , , , 1, 0)P,$$

where $P$ is an orthogonal matrix.

Apply the same orthogonal transformation to the numerator in Eq. (2):

$$\begin{aligned} P^T HWHP &= P^T HPP^T WPP^T HP \\ &= Diag(1, 1, , , , 1, 0)P^T WPDiag(1, 1, , , , 1, 0) = P^T WP_{n-1}, \end{aligned}$$

where $P^T WP_{n-1}$ denotes the submatrix of $P^T WP$ consisting of its first $(n-1)$ -th rows and columns. Hence it is also symmetric. Suppose that further diagonalizing yields

$$P^T WP_{n-1} = Q_1^T Diag(\lambda_1, \lambda_2, ..., \lambda_{n-1})Q_1, \tag{3}$$

where $\lambda_1, \lambda_2, ..., \lambda_{n-1}$ are the $n-1$ eigenvalues of the matrix $P^T WP_{n-1}$ and $Q_1$ is orthogonal. Let

$$Q = \begin{pmatrix} Q_1 & 0 \\ 0 & 1 \end{pmatrix} \tag{4}$$

$$V = PQ \tag{5}$$

Orthogonality of $P$ and $Q$ ensures that $V$ is also orthogonal. Observe that $V$ can diagonalise both $HWH$ and $H$:

$$V^T HWHV = Q^T P^T HWHPQ = Diag(\lambda_1, \lambda_2, ..., \lambda_{n-1}, 0) \tag{6}$$

$$V^T HV = Q^T Diag(1, 1, , , , 1, 0)Q = Diag(1, 1, , , , 1, 0) \tag{7}$$

Substitute Eq. (6) and (7) into (2), bivaraite Moran's $R$ can be rewritten into

$$R = \frac{n}{\sum_{i=1}^{n}\sum_{j=1}^{n} w_{i,j}} \frac{X^T V Diag(\lambda_1, \lambda_2, ..., \lambda_{n-1}, 0)V^T Y}{\sqrt{X^T V Diag(1, 1, , , , 1, 0)V^T X}\sqrt{Y^T V Diag(1, 1, , , , 1, 0)V^T Y}} \tag{8}$$

Let $V^T X = (a_1, a_2, , , , a_n)$ and $V^T Y = (b_1, b_2, , , , b_n)$. Since $V$ is orthogonal, $V^T X$ and $V^T Y$ still follows the same normal distribution, that is,

$$V^T X \sim N(0, \sigma_1^2 I) \tag{9}$$

$$V^T Y \sim N(0, \sigma_2^2 I) \tag{10}$$

or equivalently, for any $i = 1, 2, ..., n, a_i \sim N(0, \sigma_1^2), b_i \sim N(0, \sigma_2^2)$. Then $R$ can be expressed into a function of product of normal random variables:

$$R = \frac{n}{\sum_{i=1}^{n}\sum_{j=1}^{n} w_{i,j}} \frac{\sum_{i=1}^{n-1}\lambda_i a_i b_i}{\sqrt{\sum_{i=1}^{n-1} a_i^2}\sqrt{\sum_{i=1}^{n-1} b_i^2}} \tag{11}$$

Assuming that $E(R^p) = \frac{E(numerator^p)}{E(denominator^p)}$, it suffices to compute the moments of numerator and denominator respectively.
By the assumption,

$$E(a_i)E(b_i) = 0 \tag{12}$$

$$E(a_i^2)E(b_i^2) = \sigma_1^2\sigma_2^2 \tag{13}$$

$$E(a_i a_j b_i b_j) = 0 \tag{14}$$

Hence,

$$E(numerator) = \sum_{i=1}^{n-1}\lambda_i E(a_i)E(b_i) = 0 \tag{15}$$

$$E(numerator^2) = \sum_{i=1}^{n-1}\lambda_i^2 E(a_i^2)E(b_i^2) + 2\sum_{i\neq j}\lambda_i\lambda_j E(a_i a_j b_i b_j) \tag{16}$$

$$E(denominator^2) = \sum_{i=1}^{n-1} E(a_i^2)\sum_{i=1}^{n-1} E(b_i^2) = (n-1)^2\sigma_1^2\sigma_2^2 \tag{17}$$

Combining (15),(16) and (17),

$$E(R) = 0 \tag{18}$$

$$Var(R) = E(R^2) = \frac{n^2}{(\sum_{i=1}^{n}\sum_{j=1}^{n} w_{i,j})^2}\frac{\sum_{i=1}^{n-1}\lambda_i^2}{(n-1)^2} \tag{19}$$

Recall (6), it is equivalent to $HWH = VDiag(\lambda_1, \lambda_2, ..., \lambda_{n-1}, 0)V^T$. Thus, $\lambda_1, \lambda_2, ..., \lambda_{n-1}$ are also eigenvalues of $HWH$. By the property of trace and eigenvalues,

$$\sum_{i-1}^{n-1}\lambda_i^2 = tr((HWH)^2) \tag{20}$$

Substitute (20) into and represent the trace using the elements of $W$, the variance of bivariate Moran's $R$ can be expressed in the form of function of matrix $W$:

$$Var(R) = \frac{n^2\sum_{i=1}^{n}\sum_{j=1}^{n} w_{ij}w_{ji} - 2n(\sum_{i=1}^{n}(\sum_{j=1}^{n} w_{ij}\sum_{j=1}^{n} w_{ji}) + (\sum_{i=1}^{n}\sum_{j=1}^{n} w_{ij})^2}{n^2(n-1)^2} \tag{21}$$

#### 1.3 Bivariate Moran's local $R$

Local moran's $R$ at the $i$-th spot takes the form in the following equation:

$$R_i = (x_i - \bar{x})\sum_{j=1}^{n} w_{ij}(y_j - \bar{y}) + (y_i - \bar{y})\sum_{j=1}^{n} w_{ij}(x_j - \bar{x}) \tag{22}$$

Since

$$x_i - \bar{x} = \frac{n-1}{n} x_i - \frac{1}{n} \sum_{j=1, j \neq i}^{n} x_j \tag{23}$$

and each pair of $x_i$ and $x_j$ for any $i$ and $j$ are independent, for any $i$,

$$x_i - \bar{x} \sim N(0, \frac{n-1}{n} \sigma_1^2 I) \tag{24}$$

Similarly,

$$y_i - \bar{y} \sim N(0, \frac{n-1}{n} \sigma_2^2 I) \tag{25}$$

Hence,

$$E(R_i) = E(x_i - \bar{x}) \sum_{j=1}^{n} w_{ij} E(y_j - \bar{y}) + E(y_i - \bar{y}) \sum_{j=1}^{n} w_{ij} E(x_j - \bar{x}) = 0 \tag{26}$$

$$\begin{aligned} Var((x_i - \bar{x}) w_{ij} (y_j - \bar{y})) &= w_{ij}^2 (E((x_i - \bar{x})^2 (y_j - \bar{y})^2) - (E((x_i - \bar{x})(y_j - \bar{y}))^2)) \\ &= w_{ij}^2 E((x_i - \bar{x})^2 (y_j - \bar{y})^2) = w_{ij}^2 E((x_i - \bar{x})^2) E((y_j - \bar{y})^2) \\ &= w_{ij}^2 \frac{(n-1)^2}{n^2} \sigma_1^2 \sigma_2^2 \end{aligned} \tag{27}$$

$$\begin{aligned} Cov((x_i - \bar{x}) w_{ij} (y_j - \bar{y}), (y_i - \bar{y}) w_{ik} (x_k - \bar{x})) &= E((x_i - \bar{x}) w_{ij} (y_j - \bar{y})(y_i - \bar{y}) w_{ik} (x_k - \bar{x})) \\ &\quad - E((x_i - \bar{x}) w_{ij} (y_j - \bar{y})) E((y_i - \bar{y}) w_{ik} (x_k - \bar{x})) \\ &= E((x_i - \bar{x}) w_{ij} (y_j - \bar{y})(y_i - \bar{y}) w_{ik} (x_k - \bar{x})) \\ &\neq 0 \text{ if and only if i = j = k} \end{aligned} \tag{28}$$

$$\begin{aligned} Cov((x_i - \bar{x}) w_{ij} (y_j - \bar{y}), (x_i - \bar{x}) w_{ik} (y_k - \bar{y})) &= E((x_i - \bar{x}) w_{ij} (y_j - \bar{y})(x_i - \bar{x}) w_{ik} (y_k - \bar{y})) \\ &\quad - E((x_i - \bar{x}) w_{ij} (y_j - \bar{y})) E((x_i - \bar{x}) w_{ik} (y_k - \bar{y})) \\ &= 0 \end{aligned} \tag{29}$$

$$\begin{aligned} Cov((y_i - \bar{y}) w_{ij} (x_j - \bar{x}), (y_i - \bar{y}) w_{ik} (x_k - \bar{x})) &= E((y_i - \bar{y}) w_{ij} (x_j - \bar{x})(y_i - \bar{y}) w_{ik} (x_k - \bar{x})) \\ &\quad - E((y_i - \bar{y}) w_{ij} (x_j - \bar{x})) E((y_i - \bar{y}) w_{ik} (x_k - \bar{x})) \\ &= 0 \end{aligned} \tag{30}$$

Therefore,

$$\begin{aligned} Var(R_i) &= \sum_{j=1}^{n} Var((x_i - \bar{x}) w_{ij} (y_j - \bar{y})) + \sum_{j=1}^{n} Var((y_i - \bar{y}) w_{ij} (x_j - \bar{x})) \\ &\quad + 2Cov((x_i - \bar{x}) w_{ii} (y_i - \bar{y}), (y_i - \bar{y}) w_{ii} (x_i - \bar{x}) \\ &= 2 \frac{(n-1)^2}{n^2} \sigma_1^2 \sigma_2^2 \sum_{j=1}^{n} w_{ij}^2 + 2 \frac{(n-1)^2}{n^2} \sigma_1^2 \sigma_2^2 w_{ii}^2 \end{aligned} \tag{31}$$
